## supplementary figures and tables for "Adaptation to cystine limitation stress confers a targetable lipid metabolism vulnerability in pancreatic ductal adenocarcinoma"

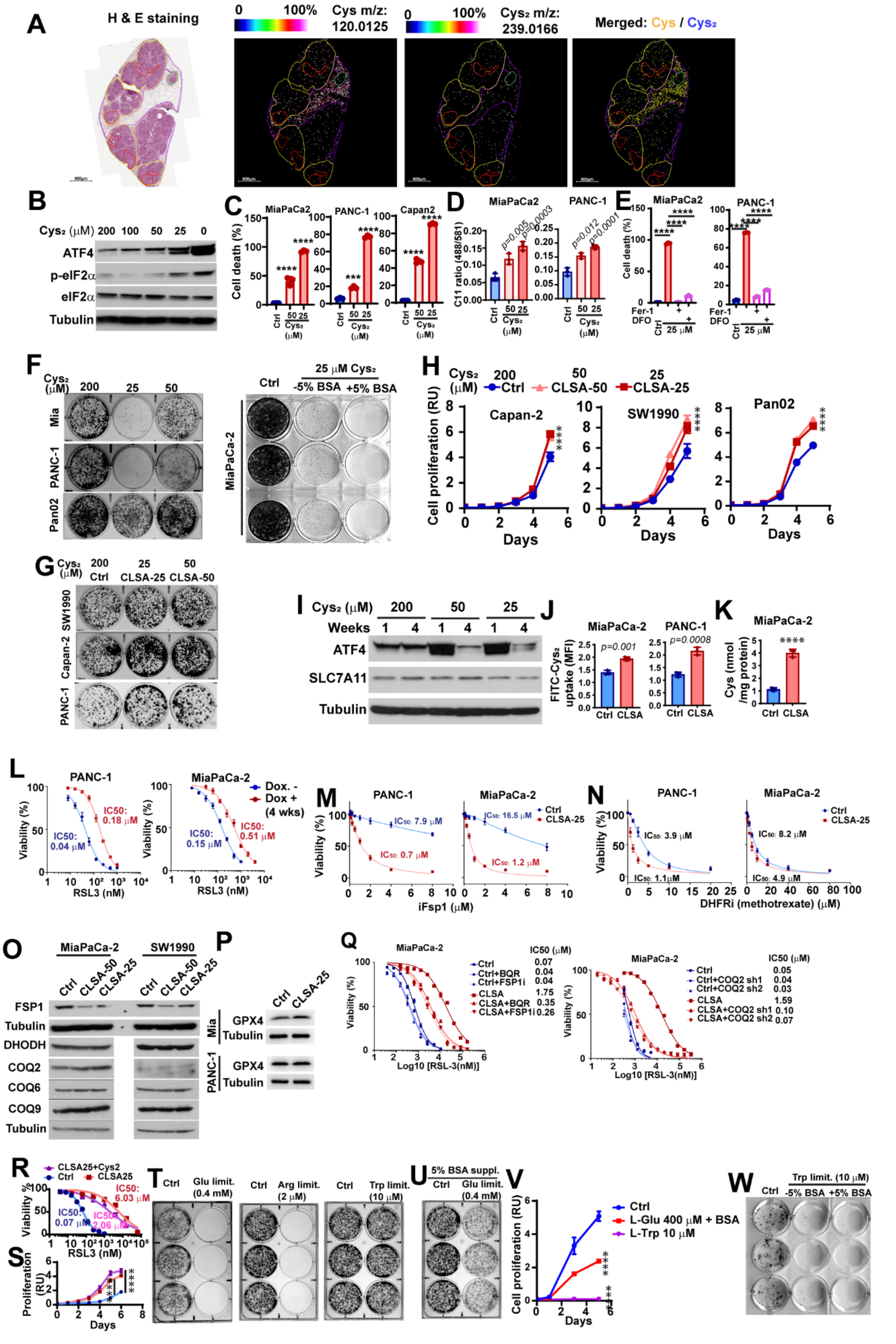

A
H & E staining
0 100% Cys m/z: 120.0125
0 100% Cys₂ m/z: 239.0166
Merged: Cys / Cys₂
B
Cys₂ (μM) 200 100 50 25 0
ATF4
p-eIF2α
eIF2α
Tubulin
C
MiaPaCa2
PANC-1
Capan2
Cell death (%)
D
MiaPaCa2
PANC-1
C11 ratio (488/581)
E
MiaPaCa2
PANC-1
Cell death (%)
Fer-1
DFO
F
Mia
PANC-1
Pan02
Ctrl
25 μM Cys₂
-5% BSA +5% BSA
MiaPaCa-2
G
200 Ctrl
25 CLSA-25
50 CLSA-50
SW1990
Capan-2
PANC-1
H
Ctrl CLSA-50 CLSA-25
Capan-2
SW1990
Pan02
Cell proliferation (RU)
Days
I
Cys₂ (μM) 200 50 25
Weeks 1 4 1 4 1 4
ATF4
SLC7A11
Tubulin
J
MiaPaCa-2
PANC-1
p=0.001
p=0.0008
FITC-Cys₂ uptake (MFI)
Ctrl CLSA
K
MiaPaCa-2
Cys (nmol /mg protein)
Ctrl CLSA
L
PANC-1
MiaPaCa-2
Dox. -
Dox + (4 wks)
IC50: 0.04 μM
IC50: 0.18 μM
IC50: 0.15 μM
IC50: 0.51 μM
Viability (%)
RSL3 (nM)
M
PANC-1
MiaPaCa-2
IC50: 7.9 μM
IC50: 0.7 μM
IC50: 16.5 μM
IC50: 1.2 μM
Ctrl
CLSA-25
iFsp1 (μM)
N
PANC-1
MiaPaCa-2
IC50: 3.9 μM
IC50: 1.1μM
IC50: 8.2 μM
IC50: 4.9 μM
DHFRi (methotrexate) (μM)
O
MiaPaCa-2
SW1990
Ctrl CLSA-50 CLSA-25
FSP1
Tubulin
DHODH
COQ2
COQ6
COQ9
Tubulin
P
Ctrl CLSA-25
Mia
PANC-1
GPX4
Tubulin
Q
MiaPaCa-2
IC50 (μM)
Ctrl 0.07
Ctrl+BQR 0.04
Ctrl+FSP1i 0.04
CLSA 1.75
CLSA+BQR 0.35
CLSA+FSP1i 0.26
Ctrl 0.05
Ctrl+COQ2 sh1 0.04
Ctrl+COQ2 sh2 0.03
CLSA 1.59
CLSA+COQ2 sh1 0.10
CLSA+COQ2 sh2 0.07
Log10 [RSL-3(nM)]
R
CLSA25+Cys2
Ctrl
CLSA25
IC50: 6.03 μM
IC50: 0.07 μM
IC50: 2.06 μM
RSL3 (nM)
S
Proliferation (RU)
Days
T
Ctrl Glu limit. (0.4 mM)
Ctrl Arg limit. (2 μM)
Ctrl Trp limit. (10 μM)
U
5% BSA suppl.
Ctrl Glu limit. (0.4 mM)
V
Ctrl
L-Glu 400 μM + BSA
L-Trp 10 μM
Cell proliferation (RU)
Days
W
Trp limit. (10 μM)
Ctrl -5% BSA +5% BSA

**Supplementary Figure 1. Adaptation to $Cys_2$ limitation stress promotes PDAC cell proliferation.**

**A,** Representative MALDI-MS images and corresponding H&E staining showing the distribution of Cys and $Cys_2$ in different regions pancreatic tissues from 8-weeks old KFC mice. Red, yellow, green and purple indicated regions of PDAC, PanIN, normal pancreatic acinar and tumor infiltrating lymphocytes, respectively.

**B,** The effects of $Cys_2$ limitation (24 hours) on the levels of ATF4, p-eIF2α in MiaPaCa-2 cells.

**C,** The effects of 72 hour CLS (50 μM or 25 μM) on PDAC cell death, as determined by PI staining and flow cytometry.

**D,** The effects of 72 hour CLS (50 μM or 25 μM) on lipid peroxidation in PDAC cells, as determined by C11-BODIPY staining.

**E,** The effects of DFO (100 μM) and ferrostatin-1 (1 μM) treatment on cell death in PDAC cells after 72-hour CLS (25 μM) treatment.

**F,** Left, the effects CLS (25 or 50 μM) on colony formation in naïve PDAC cells; right, the effect of 5% BSA supplementation on naive MiaPaCa-2 colony formation in CLS (25 μM $Cys_2$) media.

**G,** Colony formation by CLSA-25 (in 25 μM $Cys_2$), CLSA-50 (in 50 μM $Cys_2$) PDAC cells and their corresponding naïve control PDAC cells (in 200 μM $Cys_2$).

**H,** The effects of CLSA on cell proliferation in Capan-2, SW1990 and Pan02 cells.

**I,** The Western blotting analysis of the levels of ATF4 and SLC7A11 in MiaPaCa-2 cells after culture in cystine replete (200 μM) or cystine limited (25 or 50 μM) media for 1 or 4 weeks.

**J,** FITC-cystine uptake by control of CLSA25 MiaPaCa-2 and PANC-1 cells, as determined by flow cytometry.

**K,** Control or CLSA-25 MiaPaCa-2 cells were cultured in cystine replete (200 μM) media for 24 hours and the levels of intracellular cysteine were determined through LC-MS/MS after NEM derivatization.

L, The effects of chronic SLC7A11 knockdown (dox+, 4 weeks) on sensitivities to RSL-3 induced ferroptosis in PANC-1 and MiaPaCa-2 cells.

**M and N,** The sensitivities of control or CLSA-25 PANC-1 and MiaPaCa-2 cells to iFSP1 (M) or methotrexate (N) induced cell death.

**O and P,** The expression levels of FSP1, DHODH, COQ2, COQ6, COQ9 (O) and GPX4 (P) in control or CLSA PANC-1 and MiaPaCa-2 cells.

**Q,** The effects of BQR (20 μM), iFSP1 (0.1 μM) or COQ2 knockdown on sensitivities to RSL-3 induced ferroptosis in control or CLSA-25 MiaPaCa-2 cells.

**R and S,** The RSL-3 sensitivities (R) and cell proliferation (S) in MiaPaCa-2 control, CLSA-25 and CLSA-25 cells cultured in cystine replete (200 μM) media (CLSA25+$Cys_2$) for 4 weeks were determined.

**T,** The effects of glutamine limitation (400 uM), arginine limitation (2 uM) and tryptophan limitation (10 uM) on colony formation (2 weeks) in MiaPaCa-2 cells.

**U,** The effects of glutamine limitation (400 uM) and BSA supplementation (5%) on MiaPaCa-2 colony formation (2 weeks).

**V,** MiaPaCa-2 cells were cultured in conventional DMEM media or Glu-limited DMEM (0.4 mM Glu plus 5% BSA) or Trp-limited DMEM (10 uM Trp) for 4 weeks before being used for cell proliferation assay in their respective nutrient replete or nutrient-limited cell culture media.

**W,** Control MiaPaCa-2 cells or MiaPaCa-2 cells after 2-week exposure to Trp-limited (10 μM) media were harvested and used for colony formation assay in regular DMEM media or Trp-limited media (10 μM) with or without 5% BSA supplementation.

Data in (C-E) were analyzed using two sample, two-tailed paired t-test. Data in (H), (S) and (V) were analyzed using two-way ANOVA followed by Tukey's multiple comparison test. **** represents $p < 0.0001$. All error bars are mean ±SD

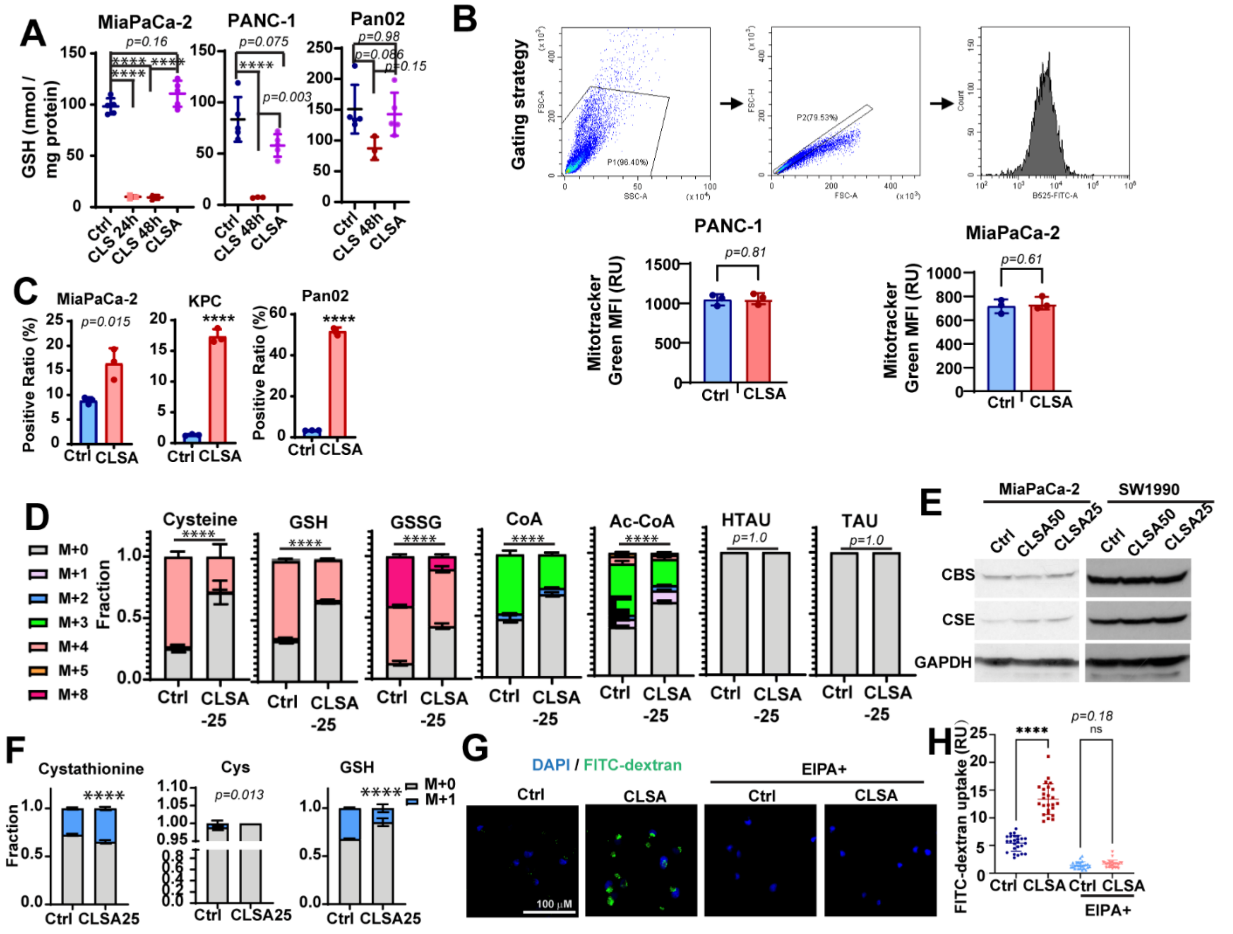

**Supplementary Figure 2. Cys metabolism in CLSA PDAC cells.**

**A,** GSH levels in PDAC cells after being cultured in $Cys_2$-limited (25 uM) media for 24-48 hours, or after adaptation to 25 uM $Cys_2$-limitation for 4 weeks were determined through LC-MS/MS.

**B,** Mitochondria volume in control or CLSA-25 PANC-1 and MiaPaCa-2 cells were determined through Mitrotracker Green staining and flow cytometry. Upper panel, gating strategy for Mitotracker Green flow cytometry. Lower panel, quantitation of flow cytometry data.

**C,** 2-NBDG assay uptake the levels of 2-NBDG positive cells in control or CLSA-25 MiaPaCa-2, KPC and Pan02 cells after 30 minute incubation with 2-NBDG.

**D,** Steady state labeling of Cys metabolites after incubation with [$^{13}C_6$, $^{15}N_2$]-$Cys_2$ for 24 hours.

**E,** The Western blotting analysis of the levels of CBS and CSE in control or CLSA MiaPaCa-2 and SW1990 cells.

**F,** Control or CLSA-25 MiaPaCa-2 cells were incubated with [1-$^{13}C$]-Serine for 4 hours and the fractional labeling of cystathionine, Cys and GSH were determined through LC-MS/MS.

**G and H**, Representative confocal microscopy images (G) and quantitation (H) showing the macropinocytosis of FITC-dextran by control or CLSA-25 MiaPaCa-2 cells with or without micropinocytosis inhibitor EIPA (25 μM).

Data in (A-D), (F) and (H) were analyzed using two sample, two-tailed, t-test. **** represents $p < 0.0001$. All error bars are mean ±SD

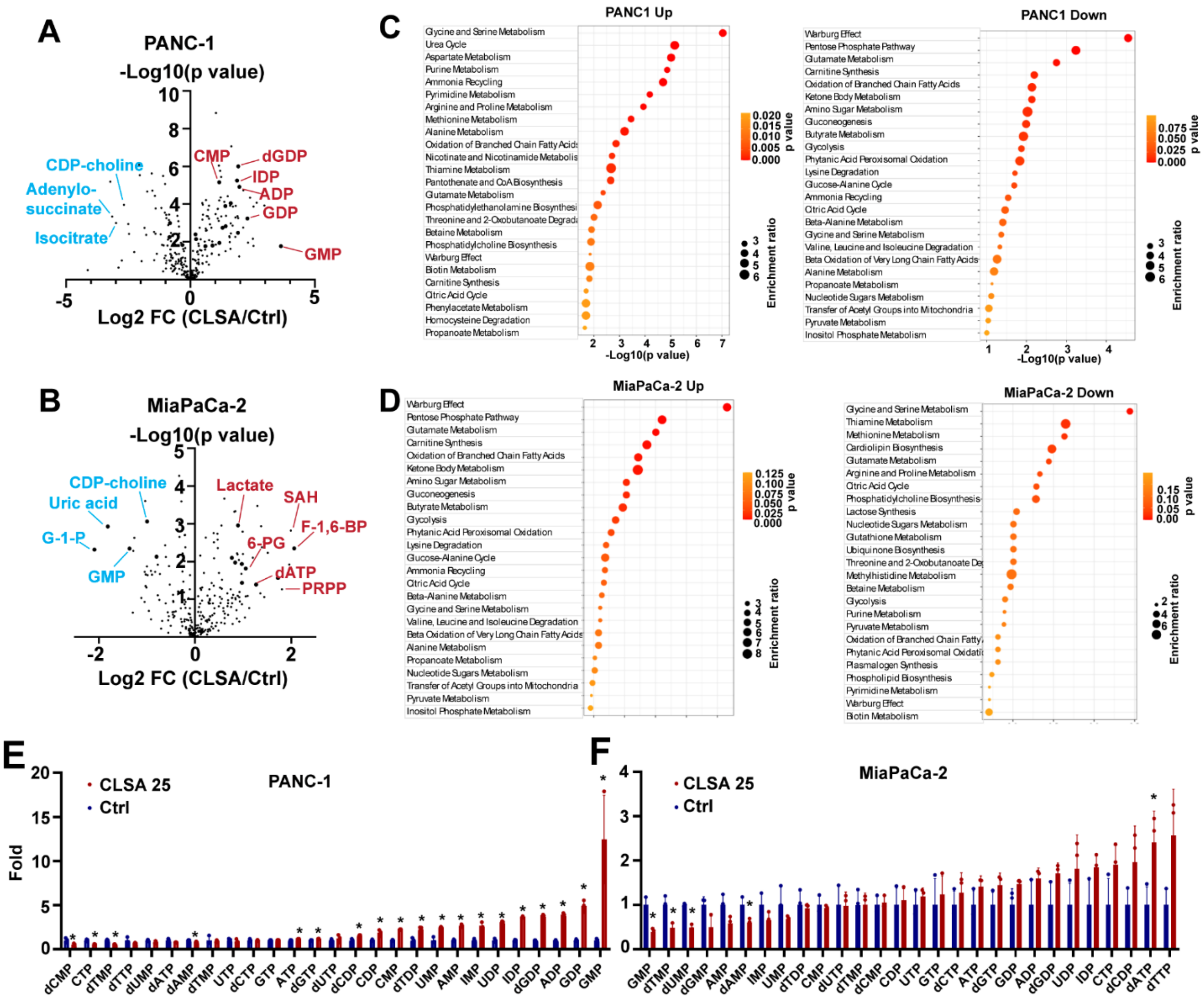

**Supplementary Figure 3. Elevated nucleotide levels in CLSA PDAC cells.**

**A and B,** Scatter dot plots showing the effect of CLSA-25 on metabolomics reprograming in PANC-1 (A) or MiaPaCa-2 (B) cells.

**C and D,** MSEA analysis of upregulated and downregulated metabolites in PANC-1 (C) and MiaPaCa-2 cells (D).

**E and F**, Bar graph showing the nucleotide levels in metabolomics screening using control or CLSA-25 PANC-1 (E) or MiaPaCa-2 (F) cells.

Data in (A), (B), (E) and (F) were analyzed using two sample, two-tailed, t-test without multiple comparison correction. * indicates $p<0.05$. All error bars are mean ±SD

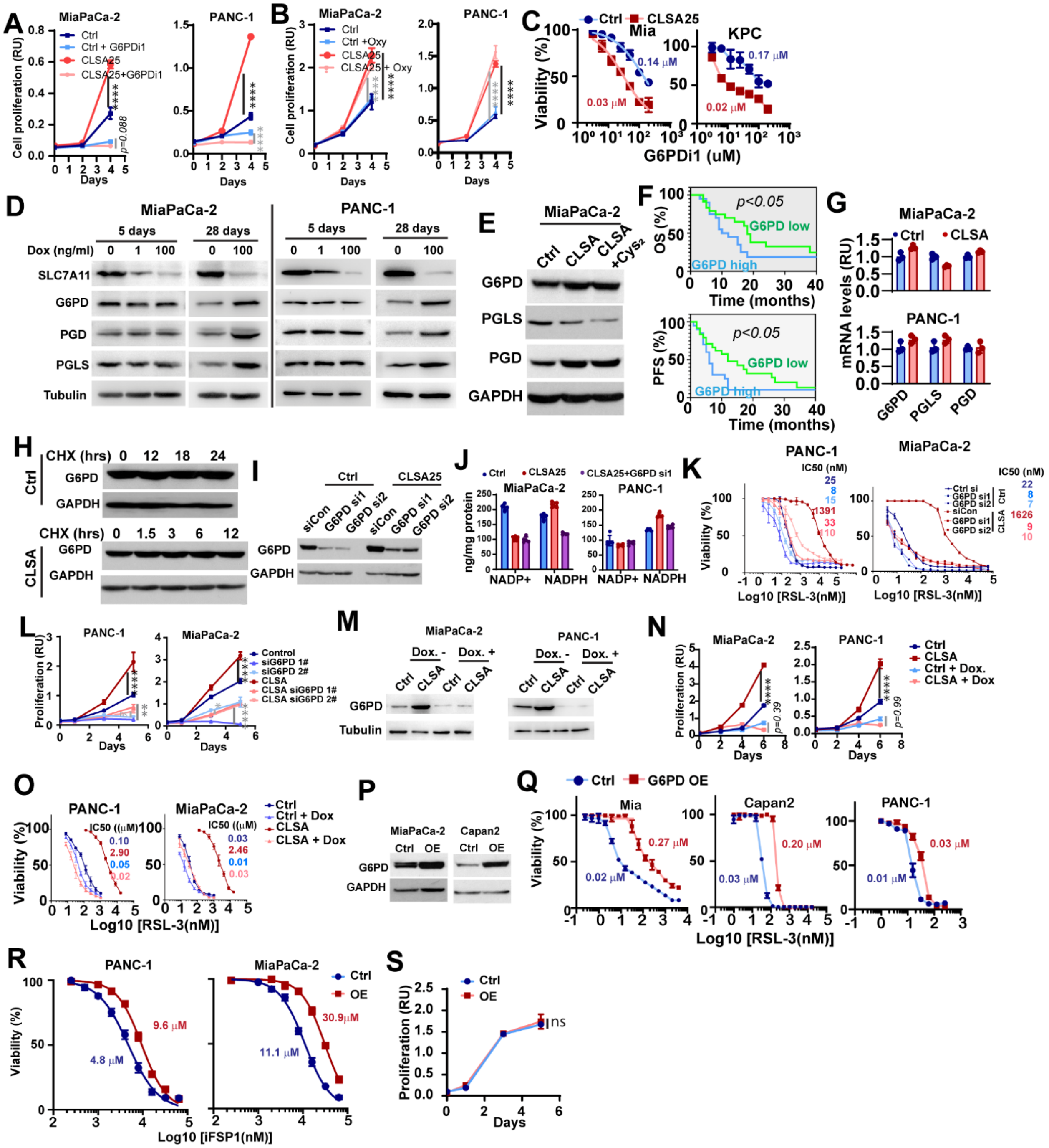

**Supplementary Figure 4. OxPPP is required for CLSA-mediated cell proliferation and ferroptosis resistance.**

**A and B**, The effects of OxPPP inhibitor G6PDi1 (50 µM) and reductive PPP inhibitor oxythiamine (50 µM) treatment on cell proliferation in control or CLSA-25 PANC-1 and MiaPaCa-2 cells.

**C,** The effect of G6PDi1 treatment on cell viability in control or CLSA-25 MiaPaCa-2 and KPC cells.

**D,** Western blotting analysis of the levels of SLC7A11, G6PD, PGLS and PGD in MiaPaCa-2 and PANC-1 cells after doxycycline-induced knockdown of SLC7A11 for 5 (left) or 28 (right) days.

**E,** Western blotting analysis of the levels of G6PD, PGLS and PGD in MiaPaCa-2 control, CLSA-25 and CLSA-25 cells cultured in cystine replete media (CLSA+$Cys_2$) for 28 days.

**F,** Kaplan-Meier analysis of the overall survival (OS) and progression free survival (PFS) of G6PD-high or G6PD-low PDAC patients (n=100).

**G,** mRNA transcript levels of G6PD, PGLS and PGD in control and CLSA-25 MiaPaCa-2 and PANC-1 cells.

**H,** The effect of cycloheximide (100 ug/mL) treatment on protein levels of G6PD in control or CLSA-25 MiaPaCa-2 cells.

**I,** The effect of G6PD siRNA1 and 2 on the protein levels of G6PD in control or CLSA-25 MiaPaCa2 cells.

**J,** NADPH and NADP+ in MiaPaCa-2 and PANC1 cells used for the calculation of NADPH/NADP+ ratios in Fig. 4J.

**K,** The effects of G6PD siRNA knockdown on sensitivities to RSL-3 induced ferroptosis in control and CLSA-25 PANC-1 and MiaPaCa-2 cells.

**L,** The effects of G6PD siRNA knockdown on cell proliferation in control and CLSA-25 PANC-1 and MiaPaCa-2 cells.

**M,** The effects of doxycycline-induced G6PD shRNA (3 Days) on the protein levels of G6PD in control or CLSA-25 MiaPaCa-2 cells.

**N and O,** The effects of doxycycline-induced G6PD knockdown on cell proliferation (N) and RSL-3 induced ferroptosis (O) in control or CLSA-25 PANC-1 and MiaPaCa-2 cells.

**P,** G6PD expression levels in PDAC cells stably expressing pLX304-vector control or pLX304-G6PD.

**Q-S,** The effects of G6PD overexpression (OE) on sensitivities to RLS-3 (Q), iFSP1 (R) or cell proliferation (S) in PDAC cells.

Data in (A), (B), (L), (N) and (S) were analyzed using two-way ANOVA followed by Tukey's multiple comparison test. **** represents $p < 0.0001$. All error bars are mean ±SD

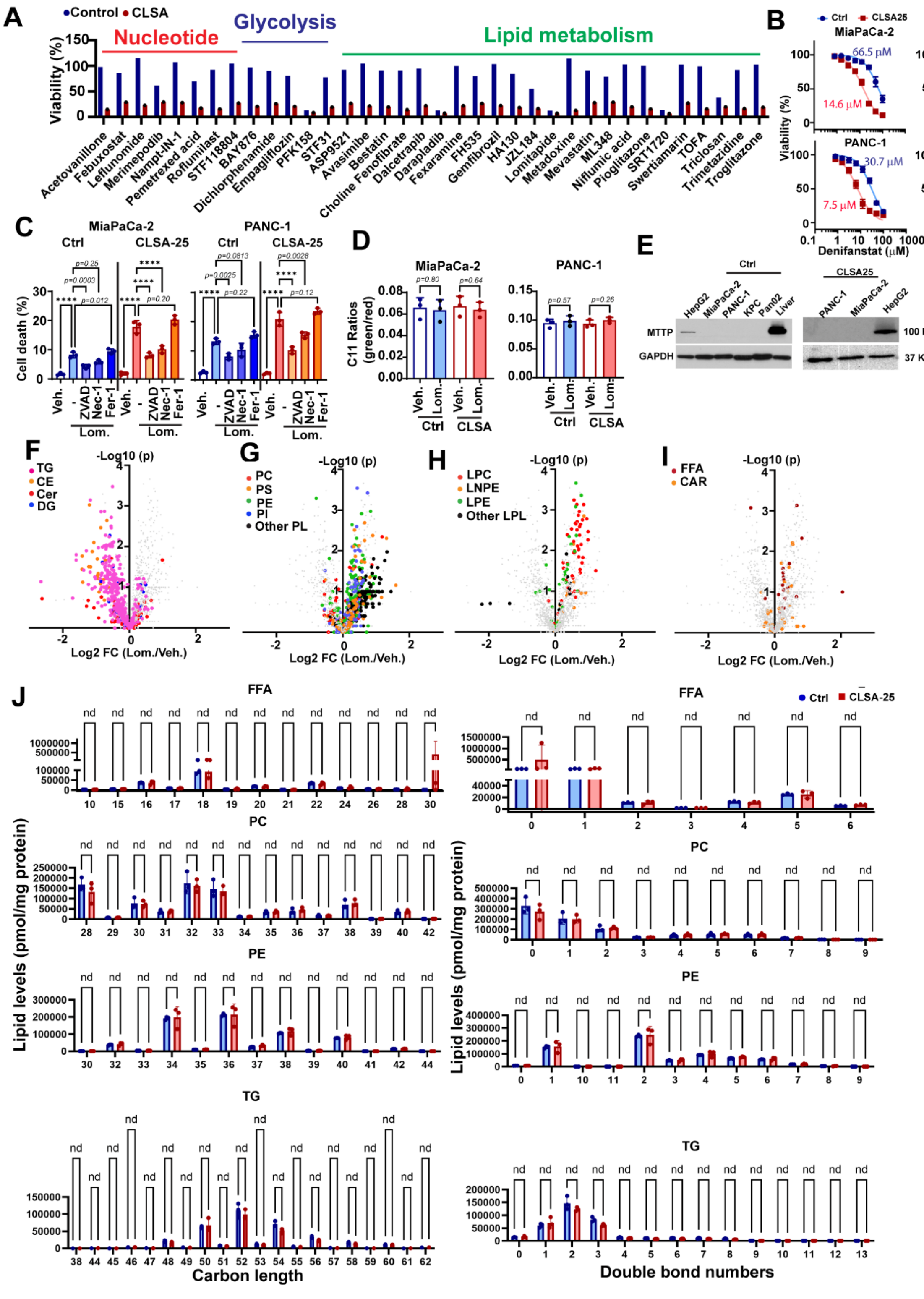

**Supplementary Figure 5. Lipid metabolism vulnerability in CLSA PDAC.**

**A,** Summary of cell viability data from positive hits in HTS drug screening.

**B,** Sensitivities of control or CLSA-25 MiaPaCa-2 and PANC-1 cells to FASN inhibitor (Denifanstat) and SCD1 inhibitor (A939572). The $IC_{50}$ for A939572 was not determined.

**C**, The effects of apoptosis inhibitor ZVAD (50 μM), necroptosis inhibitor necrostatin-1 (10 μM) and ferroptosis inhibitor ferrostatin-1 (1 μM) on lomitapide (2 μM)-induced cell death.

**D,** The effects of lomitapide treatment (2 μM, 24 hrs) on lipid peroxidation levels in control and CLSA-25 PDAC cells, as determined by C11 BODIPY fluorescence ratios.

**E,** Western blotting showing the levels of MTTP protein in naïve control or CLSA-25 human and murine PDAC cells. HepG2 and mouse liver lysate were used as positive control.

**F-I,** Volcano plots showing indicated the effects of lomitapide treatment (1 μM, 24 hrs) on changes in neutral lipids (F), phospholipids (G), lysophospholipids (H) and toxic lipid species (I) in CLSA-25 MiaPaCa-2 cells.

**J,** Analysis of the effects of lomitapide treatment on carbon length and double bond numbers in FFA, PC, PE and TG.

Data in (C) and (D) were analyzed using one way ANOVA followed by Dunnett's multiple comparison test. **** represents $p< 0.0001$. Data in (J) were analyzed using unpaired multiple t-test and * indicated false discovery rate $q<0.05$. Data in (F-I) were analyzed using two-tailed t-test without multiple comparison correction. All error bars are mean ±SD

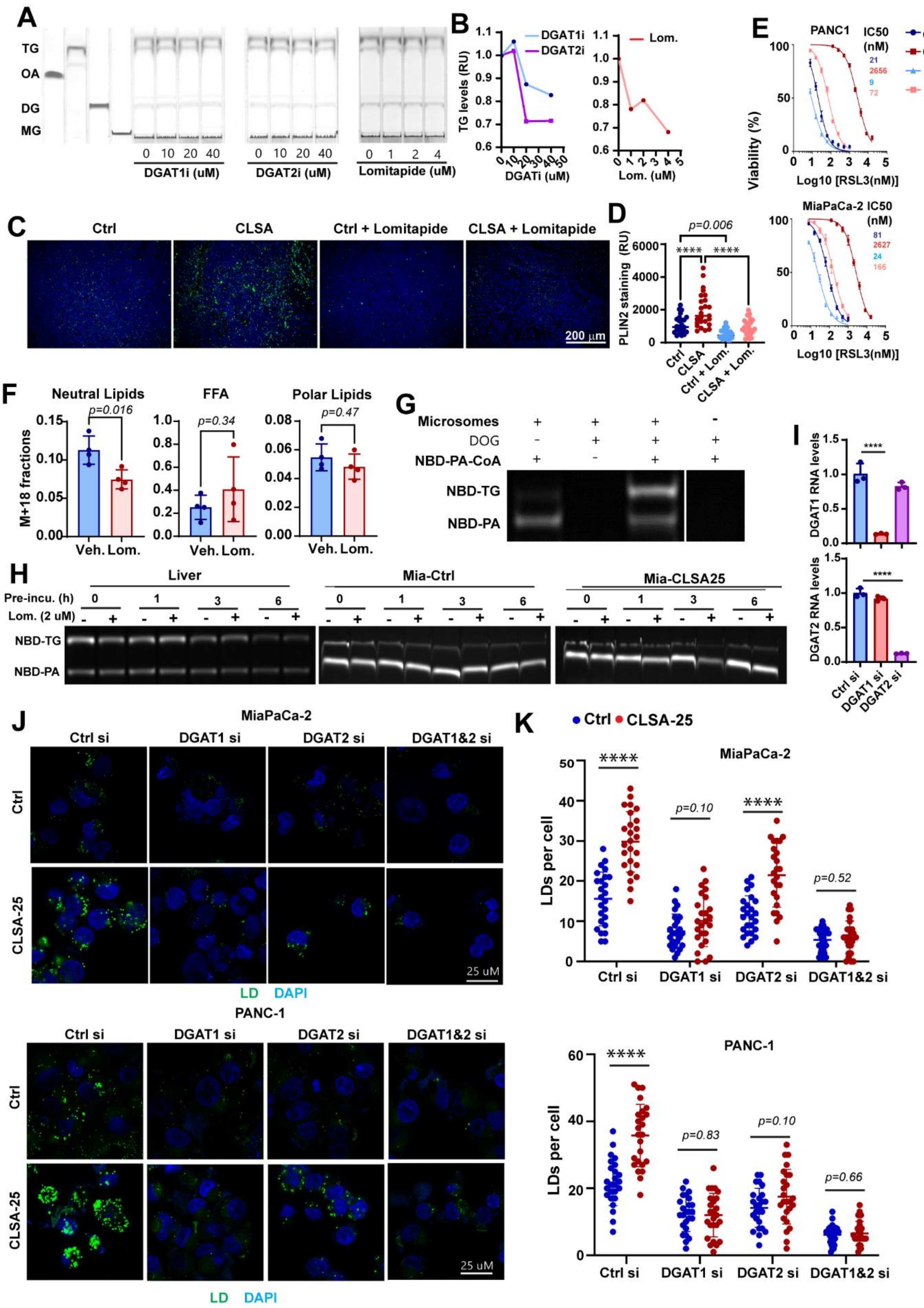

**Supplementary Figure 6. Inhibition of LD and TG synthesis by lomitapide.**

**A and B,** TLC analysis (A) and densitometry analysis (B) showing the dose-dependent effects of DGAT1i (A922500) and DGAT2i (PF-06424439) and lomitapide treatment on TG levels in MiaPaCa-2 cells.

**C and D**, Representative PLIN2 immunofluorescence staining images (C) and quantitation of staining intensities (D) of FFPE sections using xenograft tumor tissues harvested from control or CLSA-25 MiaPaCa-2 xenograft experiments with or without lomitapide treatment (10 mg/kg).

**E,** The effects of lomitapide treatment (0.5 μM) on RSL-3 induced ferroptosis in PANC-1 or MiaPaCa-2 cells.

**F,** The effects of lomitapide treatment (2 μM) on [U-$^{13}$C]-Oleate incorporation into neutral lipids, FFA and polar lipids in naive MiaPaCa-2 cells.

**G,** TLC analysis showing the synthesis of NBD-TG in the presence of DOG (200 uM), NBD-PA-CoA (25 μM) and liver microsome (5 μg protein).

**H,** The effects of pre-incubation time on the inhibitory activities of lomitapide in the *in vitro* DGAT activity assay in (G). Pre-incubation of lomitapide with microsome membrane for 1 hour or longer completely abrogated the inhibitory activities of 2 μM lomitapide.

**I,** DGAT1 siRNA and DGAT2 siRNA on the mRNA transcript levels of DGAT1 and 2 in MiaPaCa-2 and PANC-1 cells.

**J and K,** Representative confocal micrograph (J) and quantatitation (K) showing the effects of the effects of DGAT1 and DGAT2 knockdown on LD levels in control or CLSA-25 MiaPaCa-2 cells.

Data in (F), (I) and (K) were analyzed using two-tailed t-test. **** represents $p< 0.0001$. All error bars are mean ±SD

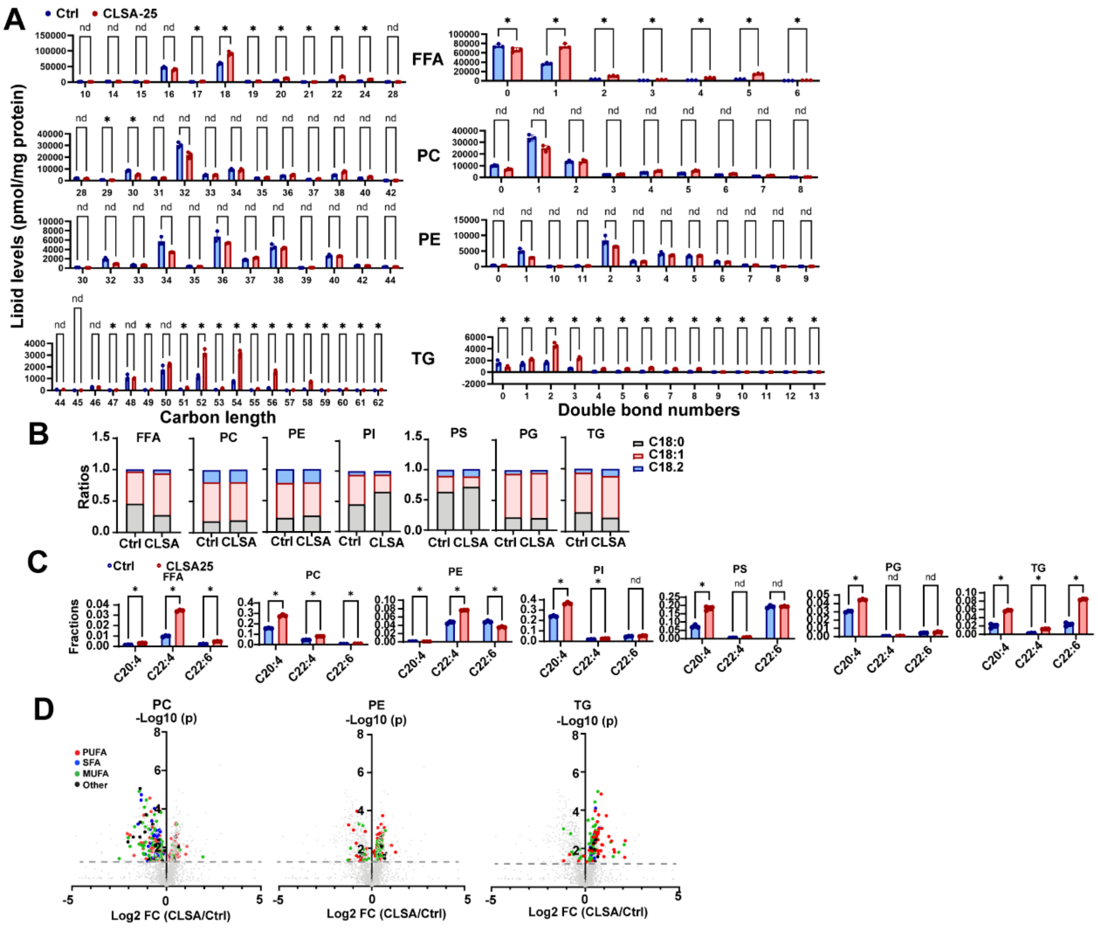

**Supplementary Figure 7. The effect of CLSA on lipidomics reprograming in PANC-1 and MiaPaCa-2 cells.**

**A,** Analysis of carbon length and double bond numbers in FFA, PC, PE and TG in control and CLSA-25 PANC-1 cells.

**B,** Analysis C18:0/C18:1/C18:2 ratios in several major lipid classes (FFA, PC, PE, PI, PS, PG and TG) in control and CLSA-25 PANC-1 cells.

**C,** The relative abundance arachidonic acid (C20:4), adrenic acid (C22:4) and DHA (C22:6) in major lipid classes (FFA, PC, PE, PI, PS, PG and TG) in control and CLSA-25 PANC-1 cells.

**D,** Scatter dot plots showing the PC, PE and TG species containing PUFA, SFA and MUFA.

Data in (A) and (C) were analyzed using unpaired multiple t-test. * indicated false discovery rate $q<0.05$. Data in (D) were analyzed using two-sided t-test without multiple comparison correction. All error bars are mean ±SD

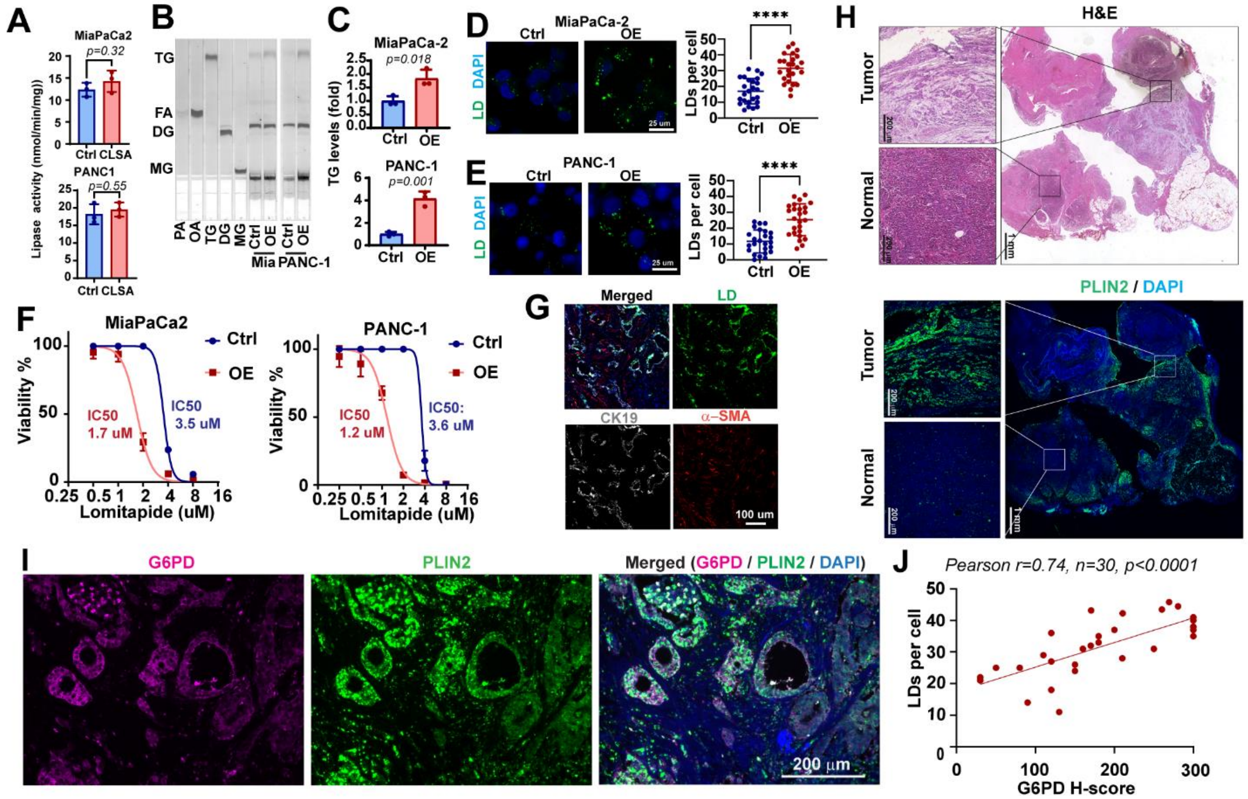

**Supplementary Figure 8. The role of G6PD in LD formation.**

**A,** Lipase activities in control or CLSA-25 MiaPaCa-2 and PANC-1 cells.

**B and C,** TLC analysis (B) and densitometry quantitation (C) showing the effects of G6PD overexpression on TG levels in MiaPaCa-2 and PANC-1 cells.

**D and E,** Representative confocal micrographs (left) and quantitation (right) showing the effects of CLSA-25 on LD levels in MiaPaCa-2 (D) and PANC-1 (E) cells.

**F,** The effects of G6PD overexpression on sensitivities to lomitapide treatment in MiaPaCa-2 and PANC-1 cells.

**G,** LD staining in tumor cells (CK19) and cancer associated fibroblasts (α-SMA) in PDAC patient tissue sections. (scale bar ??)

**H,** PLIN2 staining and H&E staining of FFPE sections from PDAC patients showing the LD levels in PDAC (tumor) and normal tissues from the pancreas of the same patient.

**I,** PLIN2 and G6PD immunofluorescence staining and representative confocal microscopy images showing the correlation between LD and G6PD expression in PDAC patient tissues.

**J,** Pearson's correlation analysis showing the correlation between G6PD and PLN2 LD staining in 30 PDAC patients.

Data in (A), (C) and (D) were analyzed using two-tailed, two sample, unpaired Student's t-test. Data in (J) were analyzed using Pearson's correlation analysis. **** represents p< 0.0001, respectively. All error bars are mean ±SD

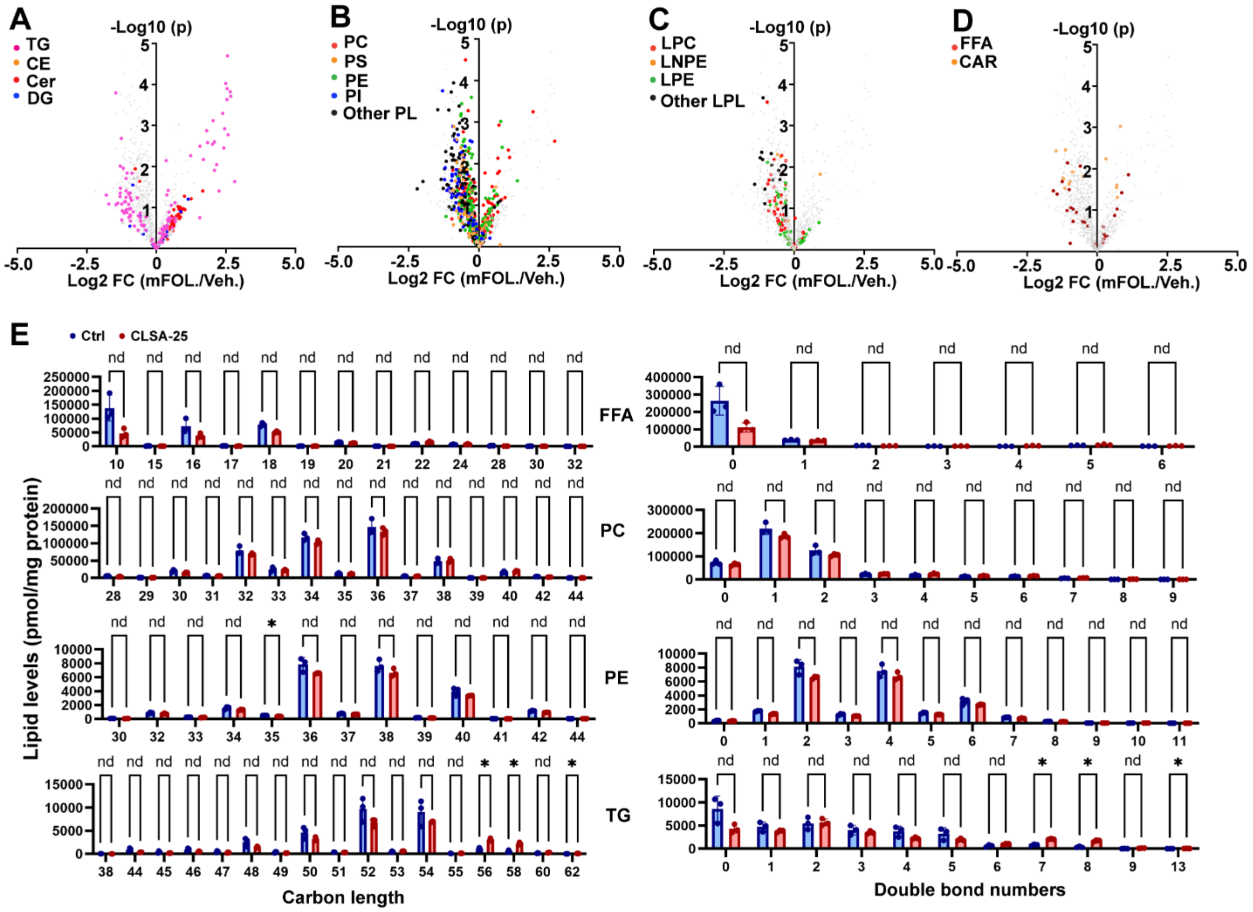

**Supplementary Figure 9. The effect of mFOLFIRINOX treatment on lipidomics changes in MiaPaCa-2 cells.**

**A-D,** Scatted dot plots showing the effects of mFOLFIRINOX treatment on neutral lipids (A), phospholipids (B), lysophospholipids (C) and toxic lipids (D) in naïve MiaPaCa-2 cells.

**E,** Analysis of carbon length and double bond numbers in FFA, PC, PE and TG in control and CLSA-25 PANC-1 cells.

Data in (E) were analyzed using unpaired multiple t-test. * indicated false discovery rate q<0.05. Data in (A-D) were analyzed using two-sided t-test without multiple comparison correction. All error bars are mean ±SD

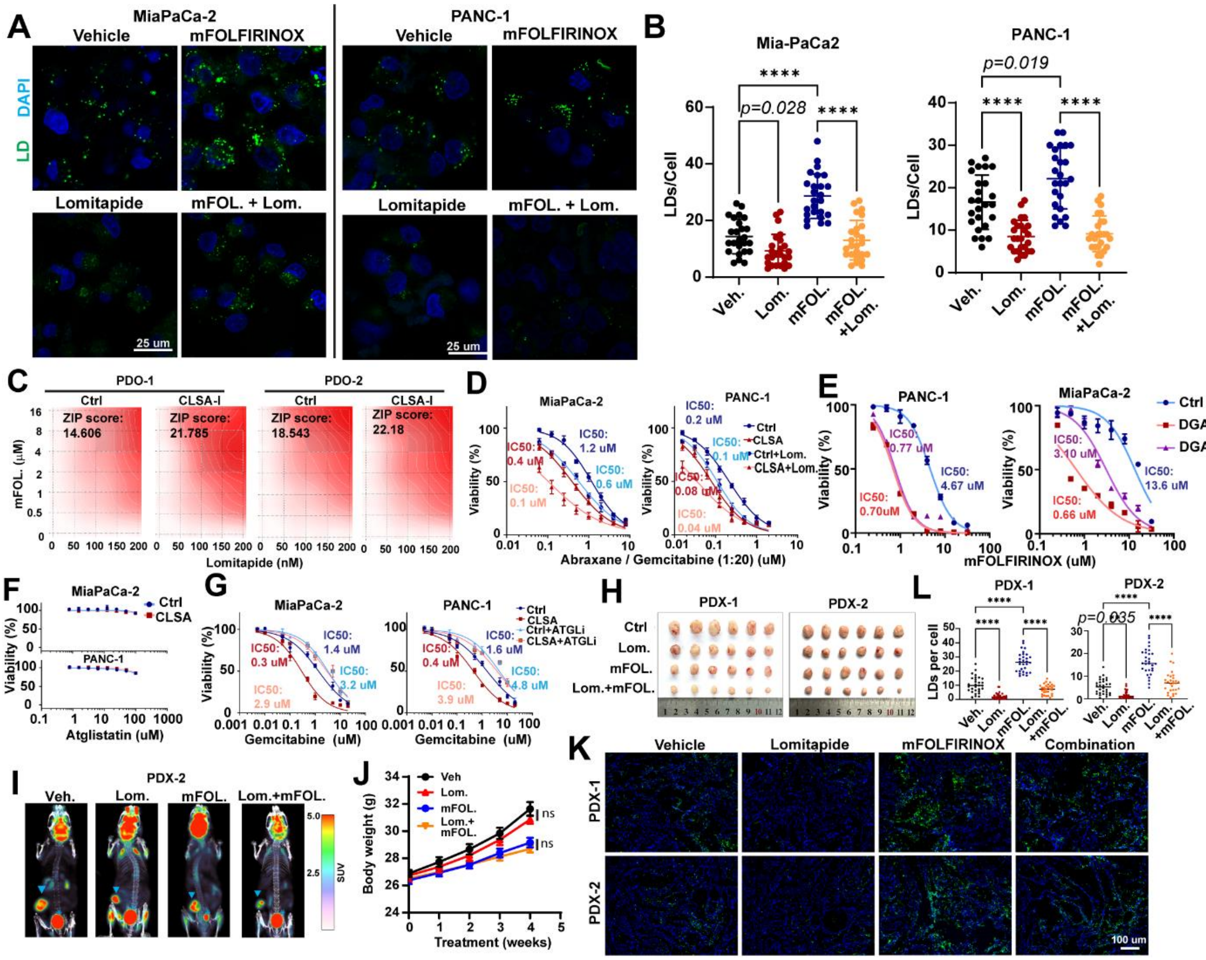

**Supplementary Figure 10. Lomitapide enhances the efficacy of chemotherapy in PDAC.**

**A and B,** Representative confocal micrographs (A) and quantitation (B) showing the effects of mFOLFIRINOX (8 µM) and lomitapide (2 µM) treatment on LD levels in MiaPaCa-2 and PANC-1 cells.

**C,** Zero Interaction Potency (ZIP) score analysis of the synergistic effects of mFOLFIRINOX and Lomitapide treatments in control and CLSA-I PDO-1 and PDO-2.

**D,** The effect of lomitapide (0.5 µM) treatment on the sensitivity of control and CLSA-25 PDAC cells to Abrexane / Gemcitabine regimen. Abrexane and Gemcitabine were added at 1:20 molar ratios and the shown concentrations are Gemcitabine concentrations.

**E,** The effects of DGAT1 and 2 knockdowns on mFOLFIRINOX sensitivities in naïve MiaPaCa-2 and PANC-1 cells.

**F and G,** The effects of ATGL inhibitor (Atglstatin) treatment alone (F) or in combination with Gemcitabine (G) on cell viability in control or CLSA-25 MiaPaCa-2 and PANC-1 cells. IC50 for (F) were not determined.

**H,** Tumors harvested from PDX-1 and PDX-2 mice treated with vehicle control, lomitapide (10 mg/kg) mFOLFIRINOX or the combination of both.

**I,** representative images of $^{18}$F-DG PET-CT scan from PDX-2 mice treated with vehicle control, lomitapide, mFOLFIRINOX or the combination of both.

**J,** bodyweight of KPC mice receiving vehicle control, lomitapide, mFOLFIRINOX or the combination of both.

**K and L,** Representative confocal micrograph (K) or quantitation (L) of LD staining of cryosections of tumor tissues harvested from PDX-1 and PDX-2 tumors treated with vehicle control, lomitapide (10 mg/kg) mFOLFIRINOX or the combination of both.

Data in (B) and (L) were analyzed using two sample, two-tailed t-test. Data in (J) were analyzed using two-way ANOVA followed by Tukey's multiple comparison test. **** represents 0.0001. All error bars are mean ±SD

**Table S1. Correlations between clinicopathological features and G6PD expression in pancreatic cancer**

| | G6PD | | | |
|---|---|---|---|---|
| | **Low** | **High** | ***p*-values** | **Spearman rank *r* value** |
| **Gender** | | | | |
| Male | 32 | 25 | 0.397 | 0.032 |
| Female | 18 | 25 | | |
| **Age** | | | | |
| > 60 | 27 | 28 | 0.755 | 0.086 |
| ≤60 | 23 | 22 | | |
| **Tumor size (cm)**[b] | | | | |
| ≤3 | 27 | 19 | 0.041 | 0.205 |
| >3 | 23 | 31 | | |
| **LN metastasis** | | | | |
| N 0 | 41 | 35 | <0.001 | 0.480 |
| N 1 | 9 | 15 | | |
| **pTNM stage** | | | | |
| I | 37 | 24 | **<0.001** | 0.376 |
| II, III | 13 | 26 | | |

Data are analyzed using Spearman's rank correlation test.

**Supplementary Table 2.** Small molecule screening data

| Category | Parameter | Description |
|---|---|---|
| Assay | Type of assay | Cell Viability / Cytotoxicity Assay (based on Cell Counting Kit-8, CCK8) |
| | Target | Cell viability of naïve PDAC cells and CLSA PDAC cells |
| | Primary measurement | Cell viability |

| | | |
|---|---|---|
| | Key reagents | Cell Counting Kit-8 test compounds DMSO (vehicle control) |
| | Assay protocol | 1. Seed target cells (naïve PDAC cells, CLSA PDAC cells) onto 96-well plates at an optimal density in their respective medium; 2. Incubate the plates at 37 °C in a 5% $CO_2$ environment for 16 hours to allow cell attachment and adaptation; 3. Add compounds from the L5200 Anti-Metabolism disease Compound Library to the wells (with DMSO as the vehicle control) to reach a final concentration of 5 μM; 4. Incubate the cells with compounds for 48 hours; 5. Measure cell viability using the Cell Counting Kit-8 according to the assay principle |
| | Additional comments | The assay was designed to investigate whether CLSA could expose exploitable metabolic vulnerabilities in PDAC cells |
| Library | Library size | 825 Compound |
| | Library composition | Anti-Metabolism disease Compound, focusing on metabolic targets including glycolysis/glucose uptake, nucleotide synthesis, and lipid metabolism |
| | Source | L5200-Anti-Metabolism disease Compound Library (TOPSCIENCE) |
| | Additional comments | The library covers multiple metabolic pathways, providing a basis for screening compounds that target metabolic vulnerabilities associated with CLSA |
| Screen | Format | 96-well plate format |

| | | |
|---|---|---|
| | Concentration(s) tested | Final concentration of 5 μM for all test compounds |
| | Plate controls | Vehicle control (DMSO) to account for non-specific effects of the solvent on cell viability |
| | Reagent/ compound dispensing system | Explorer G3 Automated Drug Screening System |
| | Detection instrument and software | EnVision 2105 Multimode Detection Instrument |
| | Assay validation/QC | Consistent incubation conditions (37 °C, 5% $CO_2$) and standardized CCK-8 assay procedures ensured reproducibility; screening focused on metabolic vulnerabilities to align with the library's anti-metabolism focus |
| | Correction factors | Not specified |
| | Normalization | Normalized to the vehicle control (DMSO) to eliminate the impact of solvent on cell viability and standardize the comparison of compound effects |
| | Additional comments | Automated compound dispensing via the Explorer G3 system guaranteed accurate and uniform delivery of test compounds |
| Post-HTS analysis | Hit criteria | Compounds that induced more than 70% inhibition of cell viability in MiaPaCa-2-CLSA cells |
| | Hit rate | ~9.2% (76 positive hits out of 825 total compounds in the library) |
| | Additional assay(s) | 1. Further functional validation of lipid metabolism-targeting hits (e.g., Lomitapide, Mevastatin, Troglitazone, Pioglitazone, Denifanstat, A939572) in naïve and CLSA PDAC cells; 2. IC50 |

| | |
|---|---|
| | determination of Lomitapide (a top hit) in CLSA PDAC cells and PDOs (IC50 range: high nanomolar to low micromolar); 3. Rescue assays using zVAD-fmk (pan-caspase inhibitor), necrostatin-1 (RIPK1 inhibitor), and ferrostatin-1 (ferroptosis inhibitor) to identify the mechanism of Lomitapide-induced cell death; 4. Detection of lipid peroxidation in Lomitapide-treated control and CLSA PDAC cells |
| Confirmation of hit purity and structure | Not specified |
| Additional comments | 1. Positive hits included glycolysis/glucose uptake inhibitors (e.g., PFK158, BAY-876), nucleotide synthesis inhibitors (e.g., Leflunomide, Pemetrexed acid), and lipid metabolism-targeting compounds (~30% of positive hits); 2. Lomitapide (a lipid metabolism-targeting hit) showed excellent efficacy in CLSA PDAC cells/PDOs, and its induced cell death involved apoptosis and necroptosis (not ferroptosis, confirmed by rescue assays and lipid peroxidation detection) |

**Table S3: Antibodies and their dilutions**

| Name | Company | Cat. No. | Dilution (Western blotting) | Dilution (IHC or IF) |
|---|---|---|---|---|
| NDUFS1 | Proteintech | 12444-1-AP | 2500 | - |
| SDHB | Santa Cruz | sc-271548 | 500 | - |
| Rieske | Santa Cruz | sc-271609 | 500 | - |

| | | | | |
|---|---|---|---|---|
| COX2 | Santa Cruz | sc-514489 | 500 | - |
| G6PD | Santa Cruz | sc-373886 | 500 | 200 |
| PGLS | Santa Cruz | sc-398833 | 250 | 200 |
| PGD | Santa Cruz | sc-398977 | 1000 | 200 |
| GCLC | Santa Cruz | sc-390811 | 500 | - |
| GCLM | Santa Cruz | sc-55586 | 500 | - |
| α-tubulin | Santa Cruz | sc-5286 | 20000 | - |
| MTTP | Santa Cruz | sc-515742 | 500 | - |
| GAPDH | Sigma | G8795 | 20000 | - |
| Histone | Millipore | 07-690 | 25000 | - |
| ATF-4 | cell Signaling | 11815S | 1000 | 200 |
| Phospho-eIF2α (Ser51) | Cell signaling | 9721S | 1000 | - |
| eIF2α Antibody | Cell signaling | 9722S | 1000 | - |
| CK19 | abcam | ab76539 | 500 | 1000 |
| α-SMA | abcam | ab7818 | 500 | 500 |
| FSP1 | santa cruz | sc-377120 | 1000 | - |
| DHODH | Proteintech | 14877-1-AP | 2000 | - |
| COQ2 | Thermo Fisher | PA5107103 | 500 | - |
| COQ6 | Santa Cruz | sc-393932 | 500 | - |
| COQ9 | Santa Cruz | sc-365073 | 1000 | - |
| GPX4 | abcam | ab252833 | 1000 | - |
| PLIN2 | abcam | ab52356 | - | 100 |
| SLC7A11 | abcam | ab307601 | 1000 | 200 |

**Table S4: Key regagents**

| Name | Company | Cat. No. |
|---|---|---|
| DMSO | sigma | D2650 |
| Fetal Bovine Serum | Atlanta Biological | S11150 |
| Penicillin/Streptomycin | Gibco | 15140163 |
| Trypsin-EDTA | Gibco | 25200-056 |
| Gibco™ DMEM, high glucose, no glutamine, no methionine, no cystine | Gibco | 21-013-024 |
| DMEM without AMINO ACID and D-Glucose | life technology | D98002710 |
| DMEM | Cytiva | SH30243.01 |
| LipidSpot™ 488 | Biotium | #70065 |
| SODIUM PALMITATE, | SIGMA ALDRICH INC | P9767 |
| Bovine Serum Albumin (FFA-free) | GoldBio | 57775 |
| RIPA Lysis and Extraction Buffer | Thermo Fisher | 88901 |
| IODINE, ROUND PARTICLES, DIAMETER 1-2.5 | SIGMA ALDRICH INC | 57655-100G-F |
| 1-MONOOLEOYL-RAC-GLYCEROL (C18:1,-CIS-9) | SIGMA ALDRICH INC | M7765 |
| OLEIC ACID BIOREAGENT, SUITABLE FOR CEL | SIGMA ALDRICH INC | O1383 |

| | | |
|---|---|---|
| 15 0 18 1 15 0 TG | FISHER SCIENTIFIC | NC2291691 |
| 16:0-18:1 DG 10MG | VWR INTL LLC | 100125-990 |
| 3-azido-7-hydroxy Coumarin | Cayman | 10596 |
| N,N-DIISOPROPYLETHYLAMINE, BIOTECH GRAD | SIGMA ALDRICH INC | 496219 |
| Oleic Acid Alkyne | Cayman | 9002078 |
| Palmitic Acid Alkyne | Cayman | 13266 |
| Collagenase from Clostridium histolytic | SIGMA ALDRICH INC | C1639 |
| HEXANE | SIGMA ALDRICH INC | 34859 |
| TETRAKIS(ACETONITRILE)COPPER(I) TETRAF | SIGMA ALDRICH INC | 677892 |
| 16-NBD-16:0 Coenzyme A | Avanti Polar Lipids | 810705P |
| 1,2-DIOLEOYL-SN-GLYCEROL, >=97%, | SIGMA ALDRICH INC | D0138 |
| L-GLUTAMINE-13C5,15N2 98+%13C-98+%15N | SIGMA ALDRICH INC | 607983 |
| D-Glucose (U-$^{13}C_6$, 99%) | Cambridge Isotope Laboratories | CLM-1396-10 |
| L-Cysteine ($^{13}C_3$, 99%; $^{15}N$, 99%) | Cambridge Isotope Laboratories | CNLM-3871-H-0.1 |
| ETHYL ETHER ANH R ACS 4L | FISHER SCIENTIFIC | E1384 |
| L-Azidohomoalanine (AHA) | Click Chemistry Tools | 1066-25 |
| Biotin-PEG4-Alkyne | Click Chemistry Tools | TA105-5 |
| AZDye 488 Alkyne | Click Chemistry Tools | 1277-1 |
| TBTA | Click Chemistry Tools | 1061-100 |
| Cycloheximide (CHX) | SIGMA ALDRICH INC | C7698 |
| DAPI (4',6-Diamidino-2-Phenylindole, Dihydrochloride) | Thermo Fisher | D1306 |
| Formalin 10% | FISHER SCIENTIFIC | SF100-4 |
| Crystal violet solution | Eng Scientific | 6100 |
| Propidium iodide | SIGMA ALDRICH INC | P4170 |
| PVDF Transfer Membranes | Millipore | IPVH00010 |
| Methanol | FISHER SCIENTIFIC | BP11054 |
| Phosphatase inhibitor cocktail | Thermo Fisher | 88667 |
| Protease Inhibitor Cocktail | SIGMA ALDRICH INC | P8340 |
| Dispase II | SIGMA ALDRICH INC | D4693 |
| L-Glutamine | Corning | 61-030-RM |
| L-Methionine | ACROS | AC166160250 |
| Sodium Pyruvate | SIGMA ALDRICH INC | P5280 |
| L-Cytine 2HCl | SIGMA ALDRICH INC | C7777 |
| D-Luciferin Sodium Salt | Regis Tech | 103404-75-7 |
| Sucrose | Millipore | 573113 |
| DNase I, RNase-free | fisher | EN0525 |
| RiboLock Rnase Inhibitor | fisher | EO0382 |
| Matrigel | Corning | CLS354248 |
| TLC plates, Silica gel 60 with concentrating zone 2.5 x 20 cm | Merk | 1118450001 |
| pENTR/D-TOPO Cloning Kit | Thermo Fisher | K240020 |
| Gateway™ LR Clonase™ II Enzyme mix | Thermo Fisher | 11791020 |

| | | |
|---|---|---|
| High-Capacity cDNA Reverse Transcription Kit | Thermo Fisher | 4368814 |
| SYBR™ Green PCR Master Mix | Thermo Fisher | 4367659 |
| Aconitase Activity Assay | Sigma | MAK051 |
| CellTiter-Glo® Luminescent Cell Viability Assay | Promega | G7572 |
| Seahorse XF Palmitate Oxidation Stress Test Kit | Agilent Technologies Inc | 103693-100 |
| Seahorse XF Cell Mito Stress Test Kit | Agilent Technologies Inc | 103015-100 |
| Seahorse XF Glycolysis Stress Test Kit | Agilent Technologies Inc | 103020-100 |
| Oasis WAX 1 cc Vac Cartridge, 30 mg Sorbent per Cartridge, 30 µm, 100/pk | Waters | 186002489 |
| Lipofectamine™ RNAiMAX Transfection Reagent | Thermo Fisher Scientific | 13778030 |
| Lactate Assay Kit | Sigma | MAK064 |
| RNeasy Kit | QIAGEN | 74106 |
| FITC-dextran | MCE | HY-128868 |
| EIPA | MCE | HY-101840 |

**Table S5: siRNA and shRNA sequence**

| | |
|---|---|
| hs.Ri.DGAT1.13.1-SEQ1 | rGrGrUrCrUrUrArCrUrGrGrUrUrGrArGrUrCrUrArUrCrACT |
| hs.Ri.DGAT1.13.1-SEQ2 | rArGrUrGrArUrArGrArCrUrCrArArCrCrArGrUrArArGrArCrCrArC |
| hs.Ri.DGAT2.13.1-SEQ1 | rArGrArUrUrCrUrGrGrArUrGrUrGrArGrGrArArGrArGrATC |
| hs.Ri.DGAT2.13.1-SEQ2 | rGrArUrCrUrCrUrUrCrCrUrCrArCrArUrCrCrArGrArArUrCrUrArG |
| hs.Ri.G6PD.13.1-SEQ1 | rGrGrUrCrArArGrGrUrGrUrUrGrArArArUrGrCrArUrCrUCA |
| hs.Ri.G6PD.13.1-SEQ2 | rUrGrArGrArUrGrCrArUrUrUrCrArArCrArCrCrUrUrGrArCrCrUrU |
| hs.Ri.G6PD.13.2-SEQ1 | rCrUrGrArCrCrUrArCrGrGrCrArArCrArGrArUrArCrArAGA |
| hs.Ri.G6PD.13.2-SEQ2 | rUrCrUrUrGrUrArUrCrUrGrUrUrGrCrCrGrUrArGrGrUrCrArGrGrU |
| SLC7A11 shRNA targeting sequence | CCTGTCACTATTTGGAGCTTT |
| G6PD shRNA targeting sequence | GTCGTCCTCTATGTGGAGAAT |
| COQ2-1 shRNA targeting sequence | CTTGGGAATTTGTGGAAAGAA |
| COQ2-2 shRNA targeting sequence | TGGGACCAGGACTATGATAAA |

**Table S6: qPCR primer sequence**

| | |
|---|---|
| hDGAT1-F | CTGGGAGCTGAGGTGC |

| | |
|---|---|
| hDGAT1-R | CACACACCAGTTCAGGATGC |
| hDGAT2-F | TCTGGGAGATGGGGCACTG |
| hDGAT2-R | GCTGCTTTTCCACCTTGGAC |
| hG6PD-F | CACCATGGCAGAGCAGGTGGCC |
| hG6PD-R | TCAGAGCTTGTGGGGGTTCACCCA |
| hPGLS-F | CCGGTTTTCGACCTGCTGAT |
| hPGLS-R | GGAGCCACAATCTTCTCCCG |
| hPGD-F | AACTCTTGGCCAAACCAGGG |
| hPGD-R | GGAGCCCATCAGGCATTGTAT |
| h.b-actin-F | CTTCGCGGGCGACGAT |
| h.b-actin-R | TAGGAATCCTTCTGACCCATGC |
| hGAPDH-F | GTCTCCTCTGACTTCAACAGCG |
| hGAPDH-R | ACCACCCTGTTGCTGTAGCCAA |
